# *finFindR*: Computer-assisted Recognition and Identification of Bottlenose Dolphin Photos in R

**DOI:** 10.1101/825661

**Authors:** Jaime W. Thompson, Victoria H. Zero, Lori H. Schwacke, Todd R. Speakman, Brian M. Quigley, Jeanine S. M. Morey, Trent L. McDonald

**Author notes:** Corresponding author: Jaime Thompson, WEST, Inc. 1610 E Reynolds St, Laramie, WY 82072.

## Abstract

1. Photographic identification is an essential research and management tool for studying population size and dynamics of common bottlenose dolphins (*Tursiops truncatus*). Photographic identification involves recognizing individuals based on unique dorsal fin markings. Manual identification of dolphins, while successful, is labor-intensive and time-consuming. To shorten processing times, we developed a series of neural networks that finds fins, assesses their unique characteristics, and matches them to an existing catalog.
2. Our software, *finFindR*, shortens photo-ID processing times by autonomously finding and isolating (*i.e.*, “cropping”) dolphin fins in raw field photographs, tracing the trailing edge of fins in cropped images, and producing a sorted list of likely identities from a catalog of known individuals. The program then presents users with the top 50 most likely matching identities, allowing users to view side-by-side image pairs and make final identity determinations.
3. During testing on two sets of novel images, *finFindR* placed the correct individual in the first position of its ordered list in 88% (238/272 and 354/400) of test cases. *finFindR* placed the correct identity among the top 10 ranked images in 94% of test cases, and among the top 50 ranked images in 97% of test cases. Hence, if a match does not exist in the first 50 images of *finFindR*’s ordered list, researchers can be almost certain (~97%) that a match does not exist in the entire catalog.
4. During a head-to-head blind test of the human-only and *finFindR-*assisted matching methods, two experienced photo-ID technicians both achieved 97% correct identification of identities when matched against a catalog containing over 2,000 known individuals. However, the manual-only technician examined 124 images on average before making a match, while the technician using *finFindR* examined only 10 images on average before finding a match.
5. We conclude that *finFindR* will facilitate equal or improved match accuracy while greatly reducing the number of examined photos. The faster matches, automated detection, and automated cropping afforded by *finFindR* will greatly reduce typical photo-ID processing times.

## Introduction

Identifying individuals from photographs is a common task in population biology, especially when research involves species that are not readily captured (IWC, 1990; Marshall & Pierce, 2012). Photo identification (photo-ID) studies can provide information on demographic rates, population size, and habitat use. In the terrestrial environment, Kelly (2001) and Sandfort (2015) applied photo identification to study cheetah and Alpine ibex. In the oceanic environment, researchers have applied photo-ID to species like whale sharks (Speed et al., 2008), sea otters (Gilkinson, Pearson, Weltz, & Davis, 2007), manatees (Langtimm et al., 2004), right whales (Hiby et al., 2013), humpback whales (Friday, Smith, Stevick, & Allen, 2000), and bottlenose dolphins (McDonald et al., 2017). Photo-ID methods recognize individuals using unique and enduring features, such as barnacle calluses on the heads of right whales or the fluke shape of humpback whales. In studies of common bottlenose dolphins (*Tursiops truncatus)*, researchers have long used the nicks, notches, and scars on dorsal fins to track the occurrence of individuals over time and to assess movements and population trends (Wells & Scott, 1990; Würsig & Jefferson, 1990; Zolman, 2002; Mazzoil, McCulloch, Defran, & Murdoch, 2004; Speakman, Lane, Schwacke, Fair, & Zolman, 2010).

Although it produces valuable results, many photo-ID methods are time-consuming and labor-intensive. When applied to bottlenose dolphins, researchers manually crop raw field photos before attempting to recognize the unique dorsal fin markings of individuals. It is common to compare images of unknown individuals to large catalogs containing thousands to tens of thousands of known individuals in order to identify a potential match. Identifying the fin in a single photo can take multiple hours, even if experts in photo-ID are familiar with the population of interest. Moreover, in some cases two separate examinations of a catalog are required to conclude a query image contains a previously unknown individual.

Software that facilitates partially automated photo-ID for bottlenose dolphins has existed for some time (Stewman, Stanley, & Allen, 1995; Auger-Méthé, Marcoux, & Whitehead, 2011; Towner, Wcisel, Reisinger, Edwards, & Jewell, 2013). Previous generations of dolphin photo-ID software generally relied on “landmarks” (anatomical reference points) to match individuals and often required substantial image processing by hand. Even after substantial processing, these systems achieve mixed accuracy and are heavily dependent on technician experience.

The rapid expansion of social media since the turn of the century has prompted improvements in photo recognition algorithms of all types. Current identification methods are typically landmark-free and generally rely on neural networks trained using machine learning methods. Image processing systems can now achieve human-level recognition rates for faces and many anthropogenic objects (Lin et al., 2014; Taigman, Yang, Ranzato, & Wolf, 2014).

We adapted social media image processing and recognition methods for application to bottlenose dolphin photo-ID tasks. Here, we introduce *finFindR*, a software system containing several neural networks that substantially shortens photo-ID processing time by autonomously cropping fins from raw photos and producing a list of likely identities sorted by likelihood. *finFindR*’s workflow generally consists of finding and isolating dorsal fins in a query (raw) image, tracing the trailing edge of fins, assigning a “score” based on distinctive characteristics, and sorting similarly “scored” identities in a catalog of known individuals by the likelihood that they match the query image. We implemented the system as an open-source R package and an associated user-friendly HTML-based application that requires no programming experience.

In this paper, we describe methods behind the general steps of *finFindR*’s workflow. As part of this work, we compared the error rates of *finFindR* to both highly experienced and novice biological technicians using a traditional manual photo-ID matching approach.

### *finFindR* workflow

*finFindR*’s workflow consists of three steps: 1) autonomous image processing to find and isolate dorsal fins in field photographs, 2) isolation of each fin’s trailing edge and computation of a “score” based on distinguishing features, and 3) computation of the proximity of an image’s “score” to the “scores” of all fins in a reference catalog. *finFindR*’s wiki (https://github.com/haimeh/finFindR/wiki) contains specific information about implementing each workflow step and should generally be considered the most up-to-date user reference for *finFindR*.

#### Step 1: Fin isolation

To autonomously identify fins in raw color (RGB) images (*e.g.*, Figure 1a), we implemented a novel neural network architecture loosely based on the “resnet” architecture (He, Zhang, Ren, & Sun, 2015). We constructed the training dataset for this network by manually labeling ~10,000 dorsal fin photographs. Manual labeling entailed outlining the fin’s edge and dolphin body by hand and assigning integer values to each region (“1” = fin edge, “2” = body; Figure 1b). Training involved passing fin photos to the network as input, allowing the network to predict regions containing fin edges and bodies, comparing predictions to labeled regions, and using backward propagation to adjust network weights. Over many training iterations, the network “learned” the characteristics of images generally associated with labels, in this case fin edges and dolphin bodies. The network outputs a pixel-based continuous value between 0 and 1 representing the likelihood that the pixel is part of a fin or body (Figure 1c). *finFindR* then creates a bounding polygon around pixels with likelihood values exceeding a sensitivity threshold. Users can specify both the sensitivity and whether extracted images should contain fins only or both fin and body. *finFindR* allows users to increase the default sensitivity threshold (0.4) to reduce the number of false fin detections. Users can also reduce the threshold to increase *finFindR*’s sensitivity for small or distant fins. Finally, *finFindR* places a rectangle around all bounding polygons in the photo and saves each to separate image files (Figure 1d).

**Figure 1:**
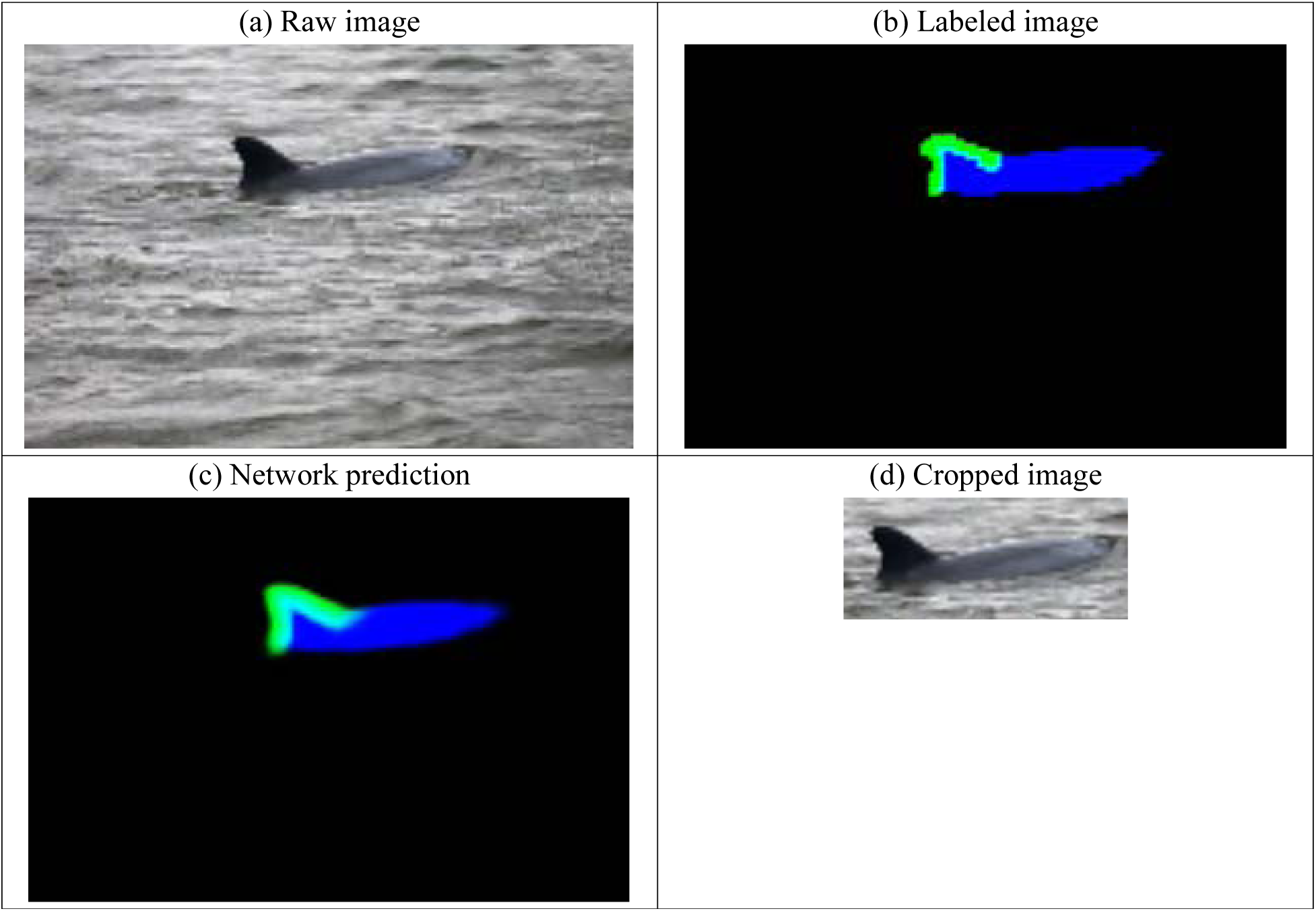
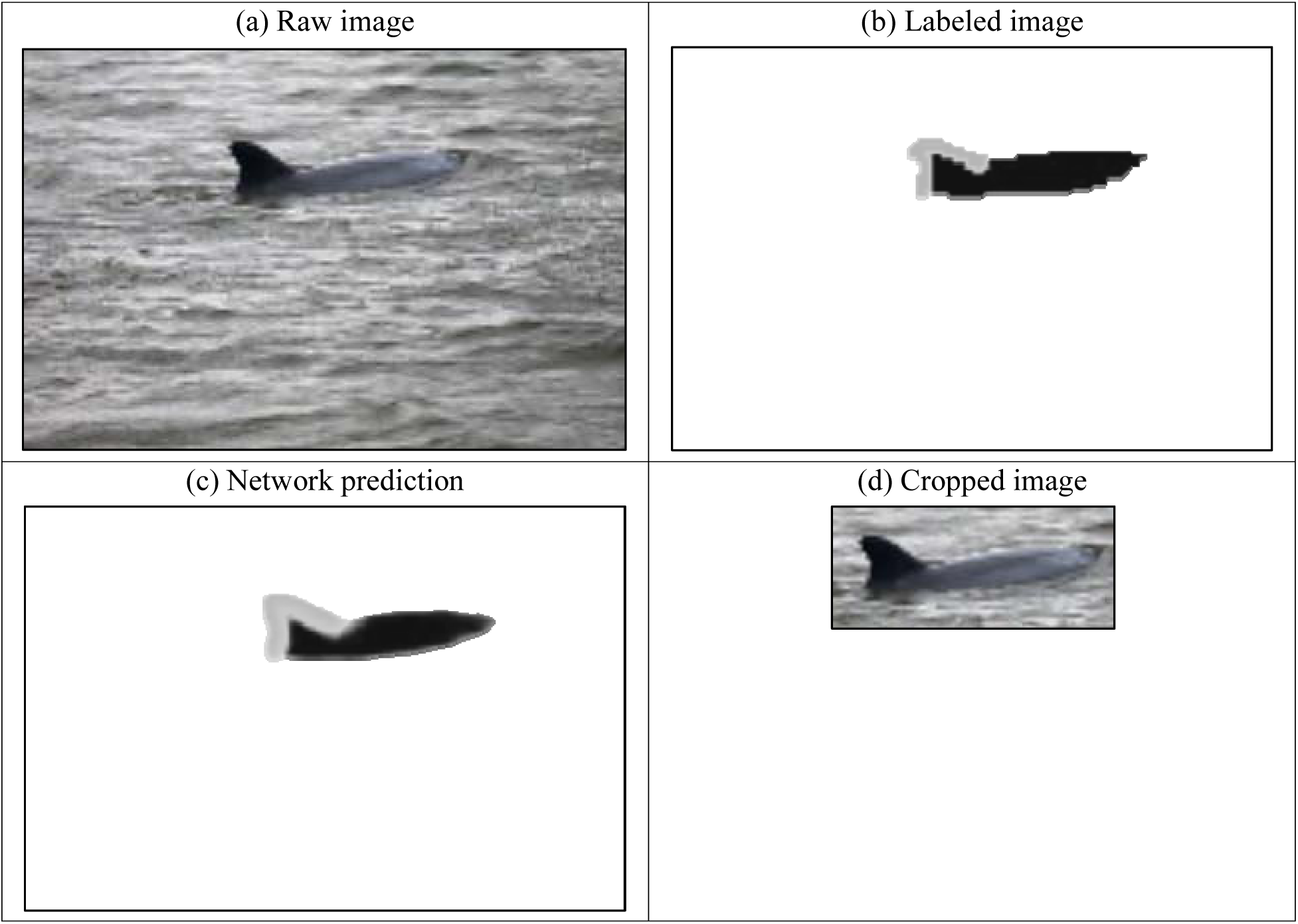
Example images illustrating fin and body isolation (Step 1 of the *finFindR* workflow). (a) The raw image; (b) manually labeled image showing location of fin edge (green) and body (blue); (c) the likelihood surface predicted by the trained network; and (d) the final cropped image.

**Figure 2:**
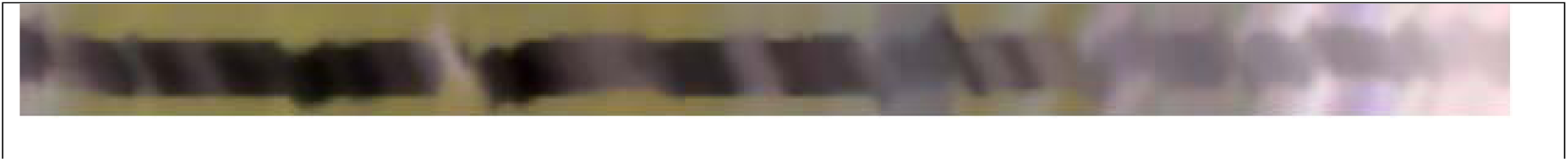
Example of a final preprocessed image input to the character extraction and mapping neural network of *finFindR*’s workflow (Step 3).

#### Step 2: Trailing edge isolation and characteristic measurement

Following fin isolation, *finFindR* isolates the trailing edge of each fin, standardizes the fin’s size, and characterizes its distinguishing features. *finFindR* isolates the trailing edge of fins using three neural networks trained to distinguish the trailing from the leading edge and to distinguish fin from body.

Once the trailing edge has been isolated, *finFindR* extracts characteristics of the trailing edge by recording red-blue-green (RGB) color values at 16 locations surrounding pixels in a large sample of pixels along the trailing edge. This sampling results in a three-dimensional matrix (hereafter, tensor) with dimensions equal to the number of pixels along trailing edge, by 16 locations, by 3 color channels. *finFindR*’s tracing tool resizes the tensor’s first dimension (*i.e.*, the fin’s trailing edge) to a standard length by applying cubic spline interpolation (Hazewinkel, 2001). Resizing the tensor in this way accommodates variable length fin edges and makes training more efficient. This standardized tensor is input to a neural network designed to distinguish individuals in the next step.

#### Step 3: Characteristic extraction and mapping

The neural network in this step is *finFindR*’s key feature and primary contribution to photo recognition technologies. The neural network in this step computes and outputs a “score” based on the fin’s distinguishing features. *finFindR* is designed to map scores to a high-dimensional mathematical space where individuals can be identified. That is, the network produces scores in a space where multiple pictures of the same fin are “close” to one another (in the high dimensional space) and “far” from the scores of other individuals. This mapping drastically reduces match-finding times when identities in the reference catalog are sorted by their proximity (“closeness”) to a query image in the high-dimensional space.

The process of mapping a tensor to high-dimensional space in a way that maximizes the distance between individuals is generally known as large-margin metric embedding (Weinberger & Saul, 2009; Faghri, Fleet, Kiros, & Fidler, 2017). We made two important modifications to make our max-margin embedding network trainable on 10,000 or fewer images. First, we induced negative curvature in the distance metric of the embedding space. This step created greater representational capacity, which ultimately allowed mapping more individuals into regions that do not already contain identities. Second, we used a squared soft-plus loss function computed on image sets containing randomly selected individuals and randomly selected photos of the same individual. Heuristically, this loss function measured distance between the embedding of a query image, those from other images of the same individual, and those known to be of other individuals.

#### Step 4: Identifying individual dolphins

To construct an ordered list of likely matches, *finFindR* computes the distance between a query image’s location in the embedding space and the location of all other images in the same space. We designed the network of Step 3 to cluster images of similar-looking fins together in the induced space in such a way that clusters of dissimilar fins largely do not overlap. For each query image, *finFindR* presents the user with both a list of the 50 “closest” identities and a hierarchical cluster of distances between individual fins. Based on these outputs, users make the final determination of matches and assign unique IDs. All vectors of characteristics (embeddings) and assigned IDs are stored in simple R objects (*i.e.*,. RData files). Users can choose to export characteristic vectors and IDs to other databases or software from R.

### Comparison and validation

Speakman et al. (2010) and Melancon et al. (2011) outline a photo matching protocol commonly used by dolphin researchers. Under this protocol, researchers first manually crop raw field images to isolate fins, then visually compare query images with those of known individuals and judge whether or not the query image matches one or more in the catalog. To assist with these tasks researchers have developed customized databases to house their images, store manually assigned characteristics, and filter large sets of images. For many years, researchers have used the *Finbase* Microsoft Access database to store, organize, and filter catalog images (Adams, Speakman, Zolman, & Schwacke, 2006). *Finbase* allows users to sort a catalog of fin images based on user-assigned attributes but does not otherwise recommend matches.

In order to evaluate the proficiency of *finFindR*’s matching algorithm, we matched a set of fin images using both the manual-only and *finFindR-*assisted methods. We compared both match agreement and the average number of inspected images required to obtain a match. Our query images consisted of 672 fin images taken during two surveys in Barataria Bay, Louisiana during May (*n* = 272 images) and September 2017 (*n* = 400 images). Of those, we easily matched 468 images based on known associates, freeze-brands, and the feature sorting capabilities in *Finbase*. Of the remaining 204 images, we identified and removed 55 duplicate photos of the same fin, leaving 149 images of unique individuals (*n* = 135 individuals from May survey; *n* = 14 individuals from September survey). We did not use any of the 672 photos during *finFindR* training.

One of us (TRS) with extensive photo-analysis experience followed the *finFindR* workflow and matched individuals among the top 50 likely matches. During this trial, *finFindR* “found a match” when it placed the correct identity of a previously seen individual among the top 50 positions of the sorted list. Another of us (BMQ) with extensive photo-analysis experience manually matched the same set of dorsal fin images using *Finbase* only. Finally, a third researcher (JSMM) with less photo-analysis experience independently repeated the manual matching process using assistance from *Finbase* only. We ensured no communication between analysts during matching. The experienced analysts checked and verified each other’s matches (TRS verified BMQ *Finbase* results, BMQ verified TRS *finFindR* results, TRS verified JSMM *Finbase* results), and conducted additional full-catalog manual searches if no match was found.

Of the 149 identities, *finFindR* failed to place 5 (3%) known individuals in the top 50 ranked identities. Assisted by *Finbase*, the other experienced analyst failed to find 6 (4%) known individuals in the catalog. The less experienced analyst failed to find 11 (7%) known individuals. While the manual and *finFindR*-assisted error rates obtained by the experienced researchers were functionally equivalent and very low, the effort required to find a match using *finFindR* was considerably less than for the manual-only method. On average, the first experienced technician examined 10 images before finding a match using *finFindR*, while the other experienced analyst examined 124 photos on average before identifying a match. In some cases, the second analyst examined well over 1000 images to find a match.

In additional, we were interested in *finFindR*’s performance on obvious matches and duplicate images. We re-tested the *finFindR* method on all images from the same surveys, not just the unique individuals (*i.e.*, all 672 images). *finFindR* achieved similar results during this trial as it did during the test of unique individuals reported above. During these latter tests, *finFindR* placed the correct identity among the top 50 ranked mages in 97% of test cases (Table 1). In addition, *finFindR* placed the correct identity in the top position during 88% of our test cases, and among the top 10 ranked images during 94% of our tests (Table 1).

**Table 1:** Accuracy of image ranks produced by *finFindR* for novel images in two sets of hold-out images. Image identities verified through full search of the image catalog by an experienced image analyst after the experiment. Here, *n* is number of images. The two sets of images reflect field image-collection bouts conducted in Barataria Bay, Louisiana.

| | May 2017 ($n = 272$) | September 2017 ($n = 400$) |
| --- | --- | --- |
| Top-ranked image was correct match | 87.50% (238/272) | 88.50% (354/400) |
| Correct match in top 10 ranked images | 94.12% (256/272) | 93.25% (373/400) |
| Correct match in top 50 ranked images | 96.69% (263/272) | 97.25% (389/400) |

## Discussion

Past software systems for identifying marine mammals made use of dolphin fin or whale fluke edge characteristics (Auger-Méthé et al. 2011; Towner et al. 2013). These programs were specifically designed for certain species and are difficult to apply to others in part because they rely on landmark features (*e.g.*, the tip of the dorsal fin) to scale the notches’ characteristics (Stewman et al., 1995). Weideman et al. (2017) used differential curvature measures in a variety of dolphin and whale fin recognition problems, but these approaches are sensitive to noise and require careful feature isolation (Stewman et al., 1995). Because dolphins can be photographed from a variety of poses and viewpoints, and hence produce slightly perturbed images of the same fin, algorithms that rely purely on angles extracted along the fluke or fin have difficulty tracing and scaling the fin. *finFindR* overcomes these limitations by extracting a series of sub-images along the trailing edge that capture features in the vicinity of the edge, including coloration of scars. Hence, *finFindR* does not depend on perfect, consistent traces of the dorsal fin to achieve its results. *finFindR* leverages information in the vicinity of the edge and is able to match a wider range of fin photos.

Based on the results of our tests, researchers can have approximately 97% confidence that matches will occur (in the top 50 images) if the query image is of a previously known individual. That is, when matches are not found using *finFindR* (not present in the top 50 ranked images), researchers can either choose to manually search the entire catalog for a match or call the image a previously unseen individual. If researchers do the latter, they can be ~97% confident that the query image does not actually occur in the catalog and that the associated image is of a new individual. If the analyses of a particular study allow lower (than 97%) accuracy, *finFindR* can be run in a fully-automated mode by associating the query image with the identity in the top slot of the ordered list. When run in fully-automated mode, researchers can expect approximately 88% correct matches.

## Conclusions

*finFindR* allows rapid and accurate comparison of dorsal fin characteristics in unprocessed photographs with those in a catalog of known individuals. *finFindR* assists researchers by sorting field photos, discarding unusable images, cropping dorsal fin images, and greatly reducing the time required to find matches. We conclude the use of *finFindR* will sustain the accuracy of experienced fin matching researchers while drastically reducing typical dolphin photo-ID processing times.

## Author’s contributions

TLM and JWT conceived of the idea and together designed various features of *finFindR*; JWT designed additional features implemented in the methodology; TRS, BMQ, JSMM, and LHS collected the photos used to train and validate *finFindR*; VHZ, JWT, and TLM led manuscript writing. All authors contributed critically to manuscript drafts and gave final approval for publication.

## Data accessibility

*finFindR* is an open-source collaboration between the National Marine Mammal Foundation (NMMF) and Western EcoSystems Technology, Inc. (WEST). The *finFindR* package and documentation are hosted at https://rdrr.io/github/haimeh/finFindR/man/finFindR-package.html.

